# Susceptibility of bacterial species commonly found in abdominal abscesses to low-dose photodynamic therapy: Effects of methylene blue concentration, fluence rate, and fluence

**DOI:** 10.1101/2024.12.18.629260

**Authors:** Darrian S. Hawryluk, Martin S. Pavelka, Timothy M. Baran

**Affiliations:** Department of Biomedical Engineering, University of Rochester, Rochester, NY; Department of Microbiology and Immunology, University of Rochester, Rochester, NY; Department of Imaging Sciences, University of Rochester, Rochester, NY

**Keywords:** Photodynamic therapy, antimicrobial, abscess, methylene blue, fluence rate, fluence

## Abstract

**Objectives:** The objective of this study was to determine the effects of methylene blue (MB) concentration, laser fluence rate, and laser fluence on efficacy of *in vitro* photodynamic therapy (PDT) for four bacteria commonly found in human abscesses.

**Materials and Methods:** PDT experiments were performed with four of the most common bacteria found in abdominal abscesses: *Escherichia coli* (*E. coli*), *Enterococcus faecalis* (*E. faecalis*), *Staphylococcus aureus* (*S. aureus*), and *Pseudomonas aeruginosa* (*P. aeruginosa*). MB concentration was varied from 50-300 µg/mL, and laser fluence rate was varied from 1-4 mW/cm^2^ at a fluence of 7.2 J/cm^2^. Higher fluence rates and fluences were explored for *P. aeruginosa*. Primary outcomes were reduction in colony forming units (CFU) following PDT, and measured MB uptake following drug incubation.

**Results:** Gram-positive bacteria (*E. faecalis* and *S. aureus*) were eradicated at all MB concentrations and laser fluence rates tested. Efficacy was reduced for *E. coli*, but still resulted in >6 log_10_ reduction in CFU when MB concentration was at least 100 µg/mL. *P. aeruginosa* required higher fluence (28.8 J/cm^2^) to achieve comparable efficacy, while increasing fluence rate did not have a significant effect on PDT efficacy. MB uptake was reduced in Gram-negative species compared to Gram-positive, particularly *P. aeruginosa*, although uptake was not significantly correlated with CFU reduction.

**Conclusions:** Gram-positive bacteria can be eradicated *in vitro* with low levels of MB (50 µg/mL), laser fluence (7.2 J/cm^2^), and laser fluence rate (1 mW/cm^2^). *E. coli* showed substantial cell killing (>6 log_10_ CFU reduction) with these same parameters. Low MB uptake and PDT efficacy in *P. aeruginosa* could be overcome by increasing the laser fluence, while increasing fluence rate did not have an effect.

## Introduction

Photodynamic therapy (PDT) is a promising antimicrobial therapy that uses light-based excitation of photosensitive drugs known as photosensitizers to generate cytotoxic reactive oxygen species^1^. PDT efficacy has been demonstrated against a wide variety of bacteria and fungi in both *in vitro* and clinical settings^2,3^. Importantly, antibiotic resistant bacteria are vulnerable to PDT, and concurrent PDT may in fact increase susceptibility to antibiotics^4^. Based on this proven success, we performed a Phase 1 clinical trial that demonstrated the safety of PDT with the photosensitizer methylene blue (MB) for treatment of human abdominopelvic abscesses. Subjects that received higher PDT fluence also showed improvements in the rate of resolution of clinical symptoms following PDT relative to those treated at lower fluence^5^.

Abscesses result from acute microbial infection, and consist of a local collection of infected fluid surrounded by a fibrous capsule^6^. While percutaneous drainage and antibiotic treatment are considered standard of care, cure rates can vary widely based on anatomical site and microbial population^7,8^. Antibiotic resistance is also a major concern, as antibiotic-resistant pathogens have been found in abscesses^5,9,10^, and the prevalence of antibiotic-resistant bacteria is increasing broadly^11^. Abscesses are a common condition, with over 28,000 abdominal drainages reimbursed by Medicare in 2013^12^ and individual U.S. hospitals treating ∼350-450 patients annually^13,14^. A wide variety of bacteria and fungi are commonly found in abscesses, with some of the most common being *Escherichia coli, Staphylococcus aureus*, and *Enterococcus faecalis*^5,15,16^. To effectively treat abscesses clinically with PDT, it is therefore important to understand the dose components that can maximize efficacy across the most common bacteria encountered.

PDT efficacy is largely a product of the combined drug-light dose, which generally consists of the photosensitizer concentration and absorbed light fluence^17^. The treatment fluence rate is also an important factor, as a minimum fluence rate is required to overcome cellular antioxidant defense and repair^18^ and oncologic PDT has shown variable efficacy as a function of fluence rate^18,19^. The PDT dose required to achieve bacterial eradication is commonly referred to as the threshold dose, which is analogous to the minimum bactericidal concentration defined for conventional antibiotics. However, this threshold dose can vary between microorganisms. For the case of abscesses, this is further complicated by the treatment geometry. While a uniform fluence rate and fluence can be ensured for *in vitro* or superficial *in vivo* illumination, the complex morphology of abscesses results in a range of fluence rates across the abscess surface for a fixed optical power delivered. Although patient-specific treatment planning can be used to improve homogeneity of the light dose^13,20,21^, it is unknown whether varying the fluence rate for a given fluence results in variable outcome for these bacterial species.

As with conventional antibiotics, differences in PDT susceptibility also exist between Gram-positive and Gram-negative bacteria^22^. For certain photosensitizers, this can be explained by a lack of penetration through the more negatively charged lipopolysaccharide outer membrane present in Gram-negatives. However, for cationic photosensitizers such as methylene blue, their positive charge results in substantial bacterial uptake^23^. It is therefore unclear whether reductions in PDT efficacy for Gram-negative bacteria are simply a result of reduced photosensitizer penetration and retention, which could be overcome by increasing drug concentration. Gramnegative bacteria also generally require higher light fluence to achieve comparable reductions in bacterial burden relative to Gram-positive bacteria^24^. As the PDT dose is a product of drug and light, it is possible that increasing the light component of the dose could compensate for reductions in photosensitizer uptake.

The strong optical absorption of methylene blue also places limitations on the maximum concentration that can be used clinically. We have found that administration of an MB concentration of 1 mg/mL delivered to abdominopelvic abscesses results in high optical absorption, even after rinsing the abscess with saline^25^. This high absorption can result in high laser power required to achieve desired fluence rate targets^20,21^, potentially increasing patient risk and reducing the number of patients that could be benefit from this therapy. It is therefore desirable to reduce the MB concentration used clinically, to reduce the laser power required and ensure patient safety.

To strengthen future clinical applications, we therefore investigated the effects of light fluence rate and administered MB concentration on PDT efficacy in four representative bacteria commonly found in abscesses. We further examined differences in MB uptake between these bacteria, to determine whether differences in PDT efficacy could be explained purely by penetration and retention of MB. Finally, we escalated both the fluence rate and fluence in PDT treatment of the least susceptible of these, *Pseudomonas aeruginosa*, to determine the relative importance of these aspects in improving PDT response.

## Materials and Methods

### Bacterial Strains Used

The four bacteria used for these experiments were a non-encapsulated mutant of *Escherichia coli* K1 (*E. coli*, laboratory collection) and a clinical isolate of *Pseudomonas aeruginosa* (*P. aeruginosa*, laboratory collection), representing Gram-negative species, and *Enterococcus faecalis* OG1RF/pC194 (*E. faecalis*, provided by Robert Quivey, University of Rochester Medical Center) and *Staphylococcus aureus* (*S. aureus*, provided by Steven Gill, University of Rochester Medical Center), representing Gram-positive species. These bacteria were chosen because they are commonly found in human abscesses^5,15^.

### Bacterial Growth

Bacteria were grown overnight on an orbital shaker at approximately 150 rpm at 37×C in Brain Heart Infusion (BHI) broth. After 17 hours, samples were centrifuged at 3000x for 10 minutes at 4×C and then rinsed with 1X Phosphate-buffered saline (PBS). This process was then repeated twice more. Bacteria were then adjusted to an optical density at 600 nm (OD_600_) of 1 in PBS.

### Methylene Blue Incubation

Bacterial samples were incubated with methylene blue (MB) concentrations of 50, 100, or 300 µg/mL (Table 1) by addition of the appropriate amount of 10 mg/mL MB (Akorn, Inc., Lake Forest, IL) to 1 mL of bacterial suspension at OD_600_=1 (MB+ samples) in 1.5 mL microcentrifuge tubes. Drug-free controls instead received an equivalent volume of PBS (MB-samples). All samples were incubated on a rotating mount for 30 minutes at room temperature. Samples were then centrifuged at 3000x for 10 minutes and rinsed with PBS as before, with this process repeated twice. At this point, samples either received PDT or were used for measurement of MB uptake, as detailed below.

**Table 1:**
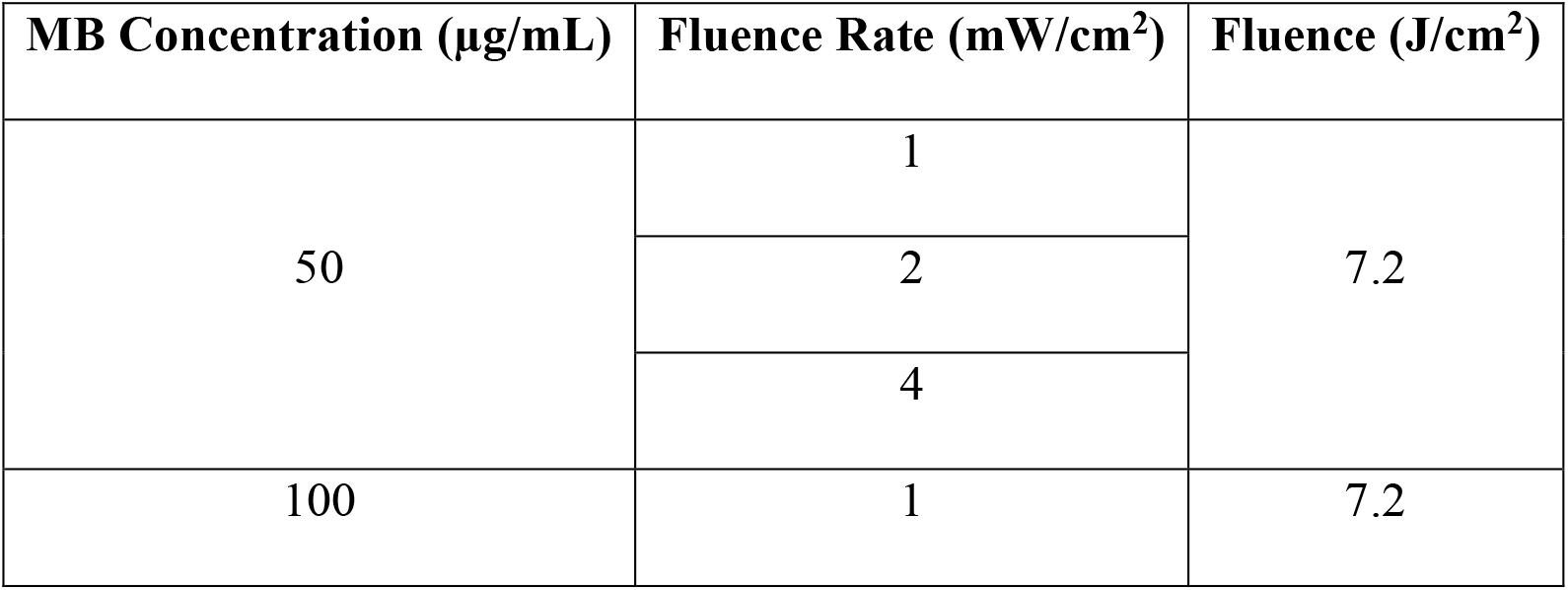
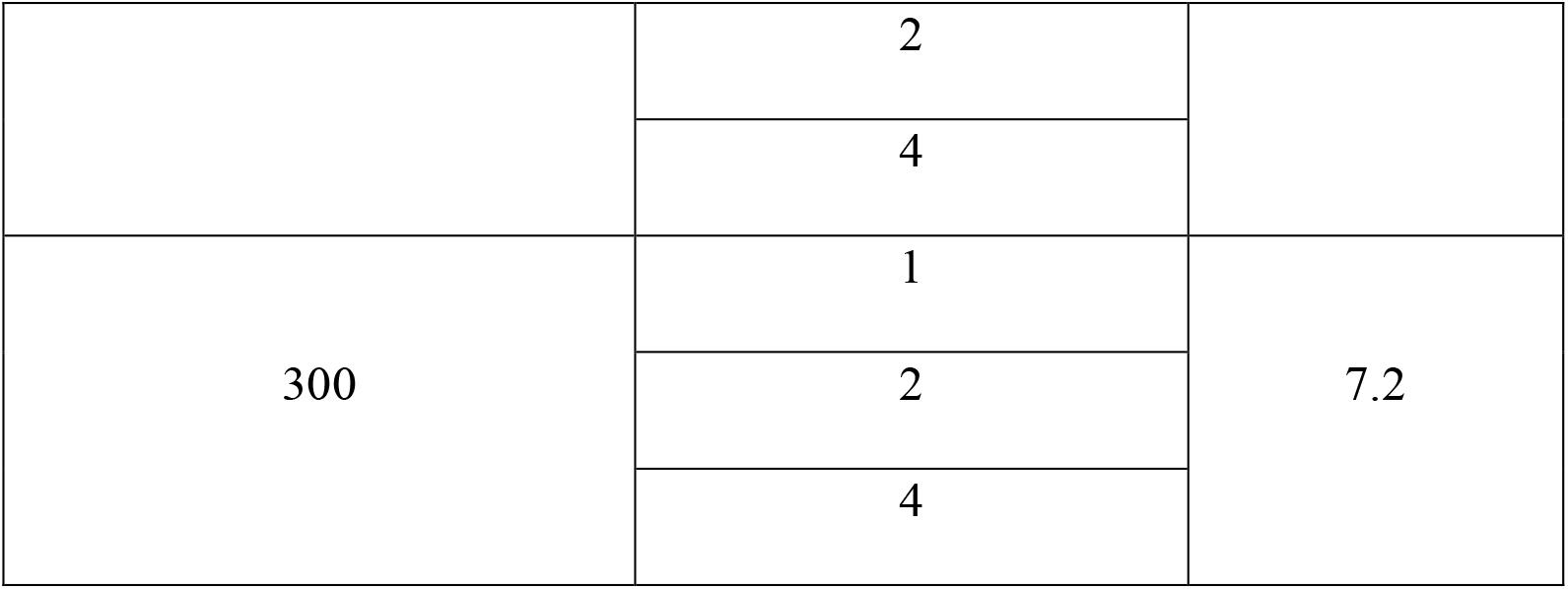
Combinations of MB concentration and light parameters used for all four bacteria.

### Photodynamic Therapy Conditions

After MB incubation, cells were resuspended in PBS in two 12-well tissue culture dishes. One of these dishes received laser treatment (L+ samples), while the other was placed under identical conditions and shielded from light (L-samples). Laser illumination was delivered by a lens-tipped optical fiber coupled to a 665 nm laser diode (LDX-3230-665, LDX Optronics, Inc., Maryville, TN). For all bacterial strains, all combinations of three MB concentrations (50, 100, or 300 µg/mL) and fluence rates of 1, 2, and 4 mW/cm^2^ were utilized, with a common total delivered fluence of 7.2 J/cm^2^ (Table 1). *P. aeruginosa* experiments also added fluence rates of 5.5 and 7 mW/cm^2^, and fluence rates of 14.4 and 28.8 J/cm^2^ (Table 2). Each experiment included a PDT-treated sample (MB+L+), a light-only control (MB-L+), a drug-only control (MB+L-), and a drug- and light-free control (MB-L-).

**Table 2:** Additional combinations of fluence rate and fluence for experiments conducted with *P. aeruginosa*.

| Fluence Rate ($\text{mW/cm}^2$) | Fluence ($\text{J/cm}^2$) |
| --- | --- |
| 4 | 7.2 |
|  | 14.4 |
|  | 28.8 |
| 5.5 | 7.2 |
|  | 14.4 |
|  | 28.8 |
| 7 | 7.2 |
|  | 14.4 |
|  | 28.8 |

Following laser illumination or control conditions, serial dilutions were plated on BHI agar plates and incubated for about 18 hours at 37×C. Bacterial colonies were counted the next day, with results reported as colony forming units per mL (CFU/mL). Results of PDT experiments are also reported as reduction in CFU/mL relative to untreated controls (MB-L-), on a log_10_ scale. Experiments were performed in triplicate, with CFU counting additionally performed in duplicate.

### Methylene Blue Uptake

Diluted samples were resuspended in PBS in 1 cm pathlength absorption cuvettes, and measured on a spectrophotometer (Cary 60, Agilent, Santa Clara, CA). Results were reported in units of optical density (OD). As measured spectra contained contributions from MB monomer absorption, MB dimer absorption, and scattering from bacteria, OD spectra were fit with a superposition of MB monomer/dimer absorption (Figure 1) and scattering. Monomer/dimer basis spectra were established based on measurement of pure MB samples. Scattering spectra were assumed to follow the form *μ*_*s*_ = *aλ*^−*b*^. This allowed for removal of the effects of cellular scattering and separation of MB monomer and dimer species.

**Figure 1:**
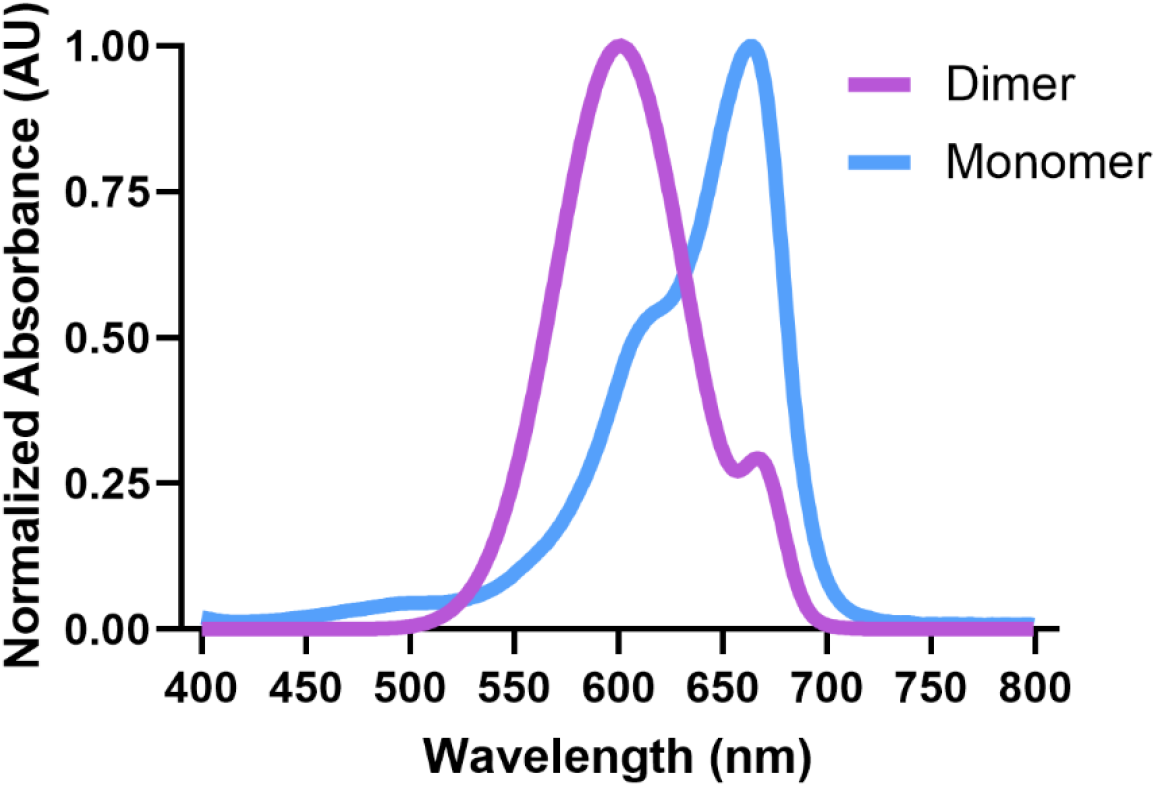
Methylene blue monomer and dimer basis functions, reported on a normalized scale

### Statistical Analysis

Differences in reduction in CFU/mL between PDT treatment conditions were analyzed using two-way ANOVA with MB concentration and light fluence rate as the main effects. Multiple comparisons were performed with Tukey’s test. The Wilcoxon test was used to compare MB uptake between bacteria. Reported correlations use the Pearson correlation coefficient. P values <0.05 were considered significant. All statistical analyses were performed in GraphPad Prism (version 10.2, GraphPad Software, LLC, Boston, MA) and MATLAB (R2023a) The Mathworks, Inc., Natick, MA).

## Results

### PDT Efficacy Across Species

Across the four species that were treated with the same parameters (Table 1), the Gram-positive bacteria were eradicated (Figure 2b and 2d). On the other hand, Gram-negative bacteria showed reduced efficacy for the same treatment parameters (Figure 2a and 2c).

**Figure 2:**
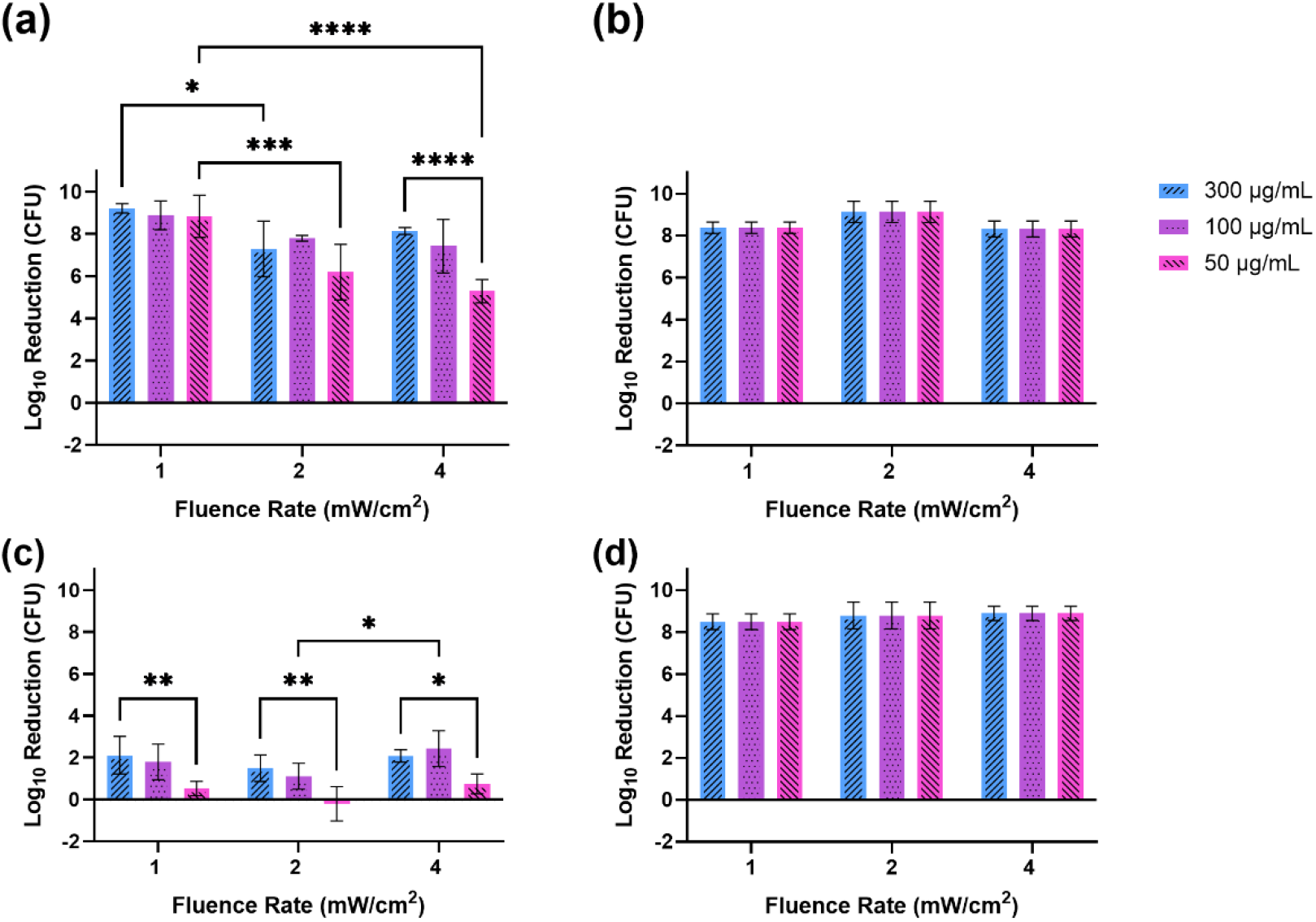
Log_10_ reduction in colony forming units (CFU)/mL as a function of fluence rate (mW/cm^2^) and MB concentration (µg/mL) used for PDT across species for **(a)** *E. coli*, **(b)** *S. aureu*s, **(c)** *P. aerugino*sa, and **(d)** *E. faecalis*. A fluence of 7.2 J/cm^2^ was used for all experiments. Solid bars correspond to mean values across experiments, while error bars correspond to standard deviation across experiments. * p<0.05, ** p<0.01, *** p<0.001, **** p<0.0001.

For *E. coli*, log_10_ reduction was significantly reduced at higher MB concentrations relative to 50 µg/mL for a fluence rate of 4 mW/cm^2^ (300 µg/mL: p<0.0001, 100 µg/mL: p=0.0036), but did not vary significantly as a function of MB concentration for other fluence rates. Increased reduction was observed at a fluence rate of 1 mW/cm^2^ for MB concentrations of 300 µg/mL (2 mW/cm^2^: p=0.012) and 50 µg/mL (2 mW/cm^2^: p=0.0002, 4 mW/cm^2^: p<0.0001), but was not a significant factor for other *E. coli* experiments.

In the case of *P. aeruginosa*, log_10_ reduction was significantly increased for 300 µg/mL MB relative to 50 µg/mL for all fluence rates (1 mW/cm^2^: p=0.0055, 2 mW/cm^2^: p=0.0024, 4 mW/cm^2^: p=0.035). At a MB concentration of 100 µg/mL, a fluence rate of 4 mW/cm^2^ resulted in significantly greater reduction relative to 2 mW/cm^2^ (p=0.038) for *P. aeruginosa* but was not significant for other fluence rate comparisons. Reduction in CFU was much smaller (p=0.0039) for *P. aeruginosa* (Figure 2c) than *E. coli* (Figure 2a). We therefore performed additional escalation of light fluence rate and fluence for *P. aeruginosa*, as described below.

### Methylene Blue Uptake

Methylene blue uptake showed similar trends across administered MB concentration and incubation time for the four species tested (Figure 3). In general, MB uptake and retention increased with MB concentration. However, measured MB concentration seemed to plateau for both incubation time and administered MB concentration beyond 20 minutes and 100 µg/mL, respectively. Amongst Gram-positive strains, MB uptake was greater for *E. faecalis* than *S. aureus* across administered MB concentrations and incubation times (p=0.0078, Figures 3b and 3d). MB uptake was significantly reduced for Gram-negative relative to Gram-positive (p<0.0001), with this reduced uptake driven by lower measured concentrations in *P. aeruginosa. P. aeruginosa* uptake was also significantly reduced relative to *E. coli* (p=0.0078, Figure 3c).

**Figure 3:**
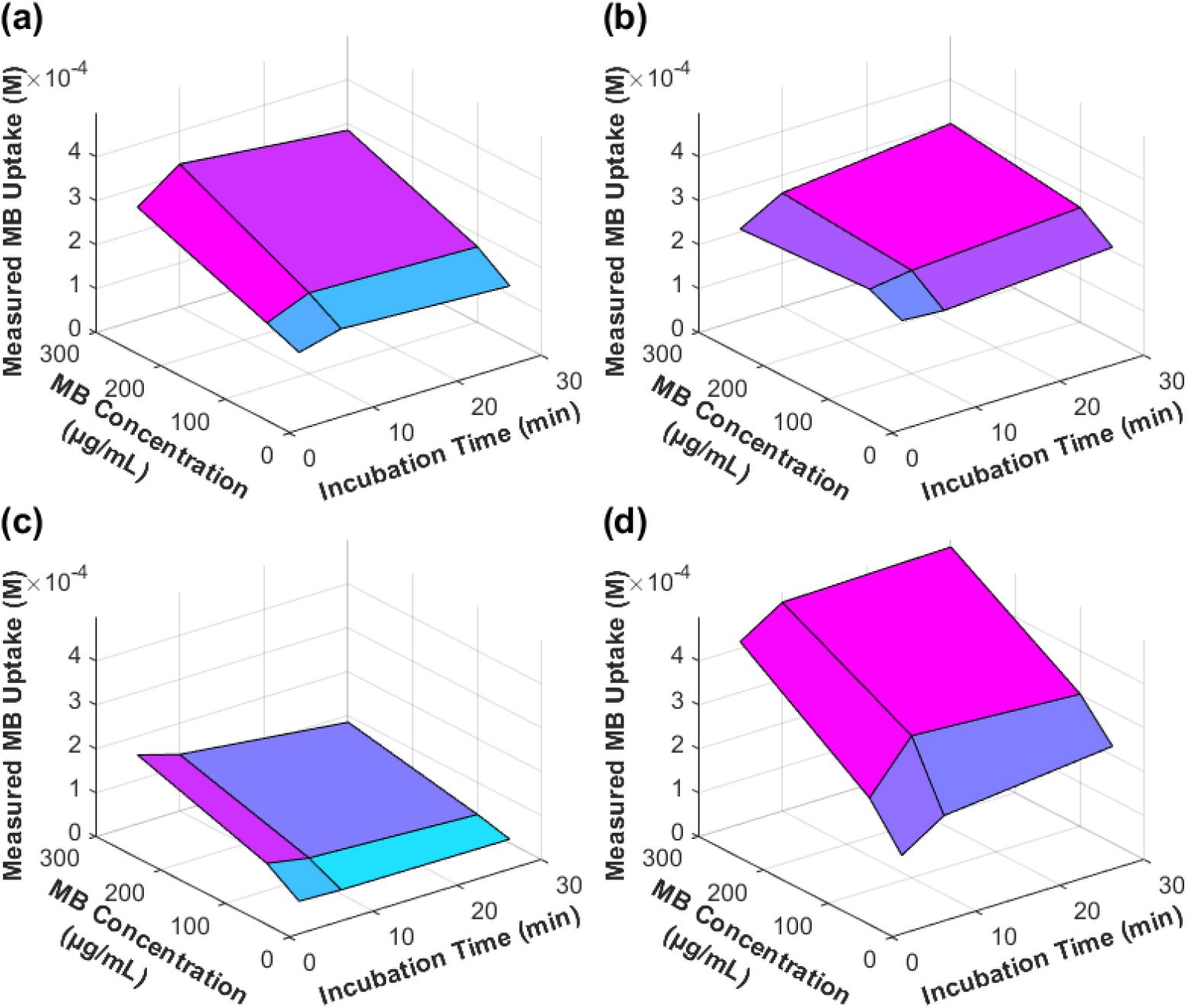
Methylene blue (MB) uptake across species as a function of administered MB concentration and incubation time for **(a)** *E. coli*, **(b)** *S. aureus*, **(c)** *P. aeruginosa*, and **(d)** *E. faecalis*. Error bars are excluded for clarity but are included in summary values in Figure 4.

When comparing log_10_ reduction in CFU to MB uptake, the Gram-positives did not show any correlation between uptake and kill, as full eradication was achieved under all conditions for these strains. Gram-negatives showed an overall positive correlation between MB uptake and reduction in CFU, which also seemed to plateau when administered MB concentration increased from 100 µg/mL to 300 µg/mL (Figure 4a-c). However, these correlations were not significant for any fluence rates examined.

**Figure 4:**
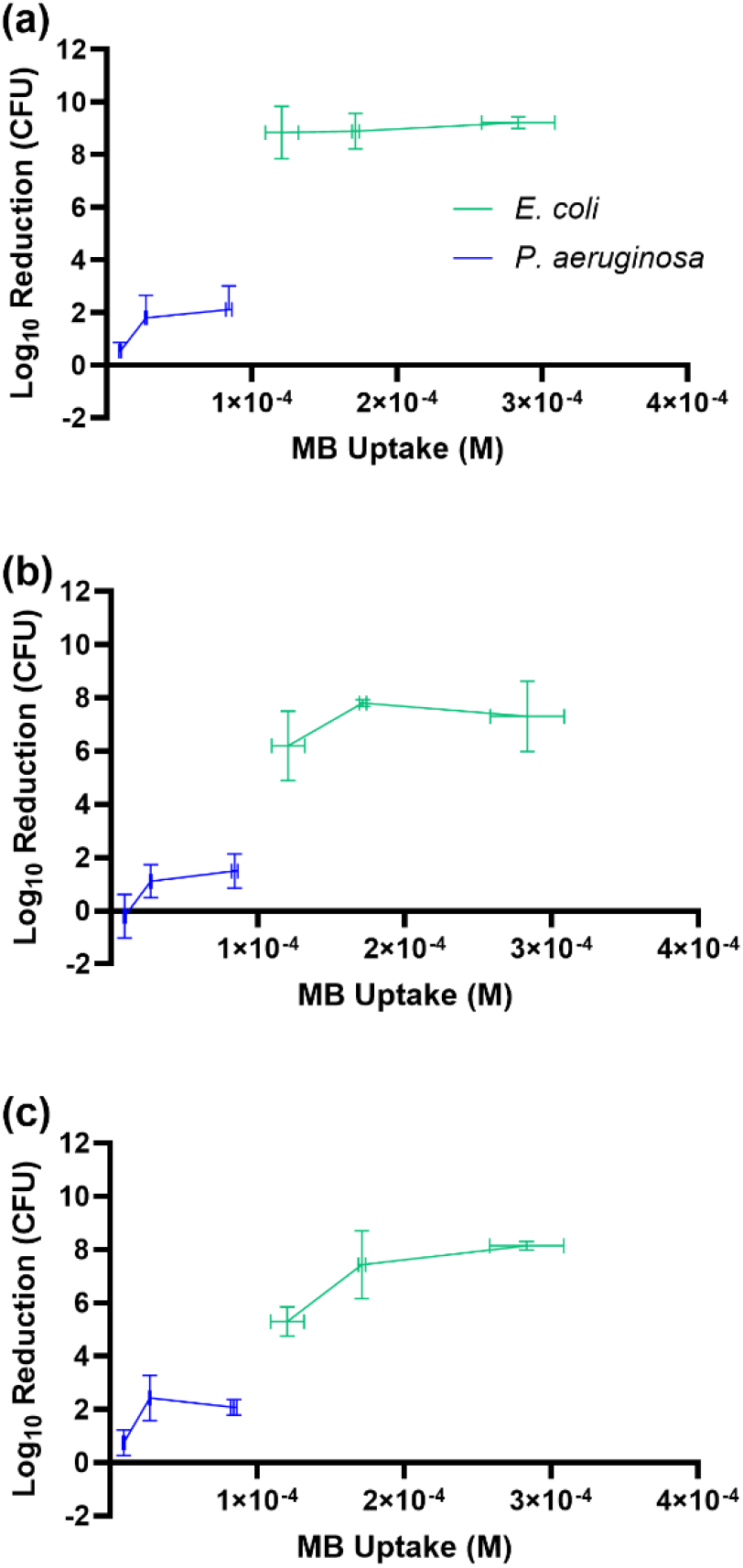
Correlation between log_10_ reduction in CFU and MB uptake in both *E. coli* and *P. aeruginosa* for the fluence rates **(a)** 1 mW/cm^2^, **(b)** 2 mW/cm^2^, and **(c)** 4 mW/cm^2^. Data points correspond to mean values across replicates, with horizontal error bars corresponding to standard deviation in measured MB uptake, and vertical error bars corresponding to standard deviation in log_10_ reduction in CFU.

#### Effects of light fluence rate and fluence for P. aeruginosa

The effects of fluence, fluence rate, and MB concentration on log_10_ reduction in CFU for *P. aeruginosa* are shown in Figure 5. Increasing the fluence rate did not result in any significant differences (p>0.05 in all cases). However, increasing the fluence resulted in significant improvement in antimicrobial effect (p<0.05 in all cases). For example, at a fluence rate of 4 mW/cm^2^, there was significant difference between the fluences of 7.2 J/cm^2^ and 28.8 J/cm^2^ for all three MB concentrations (p<0.0001 in all cases).

**Figure 5:**
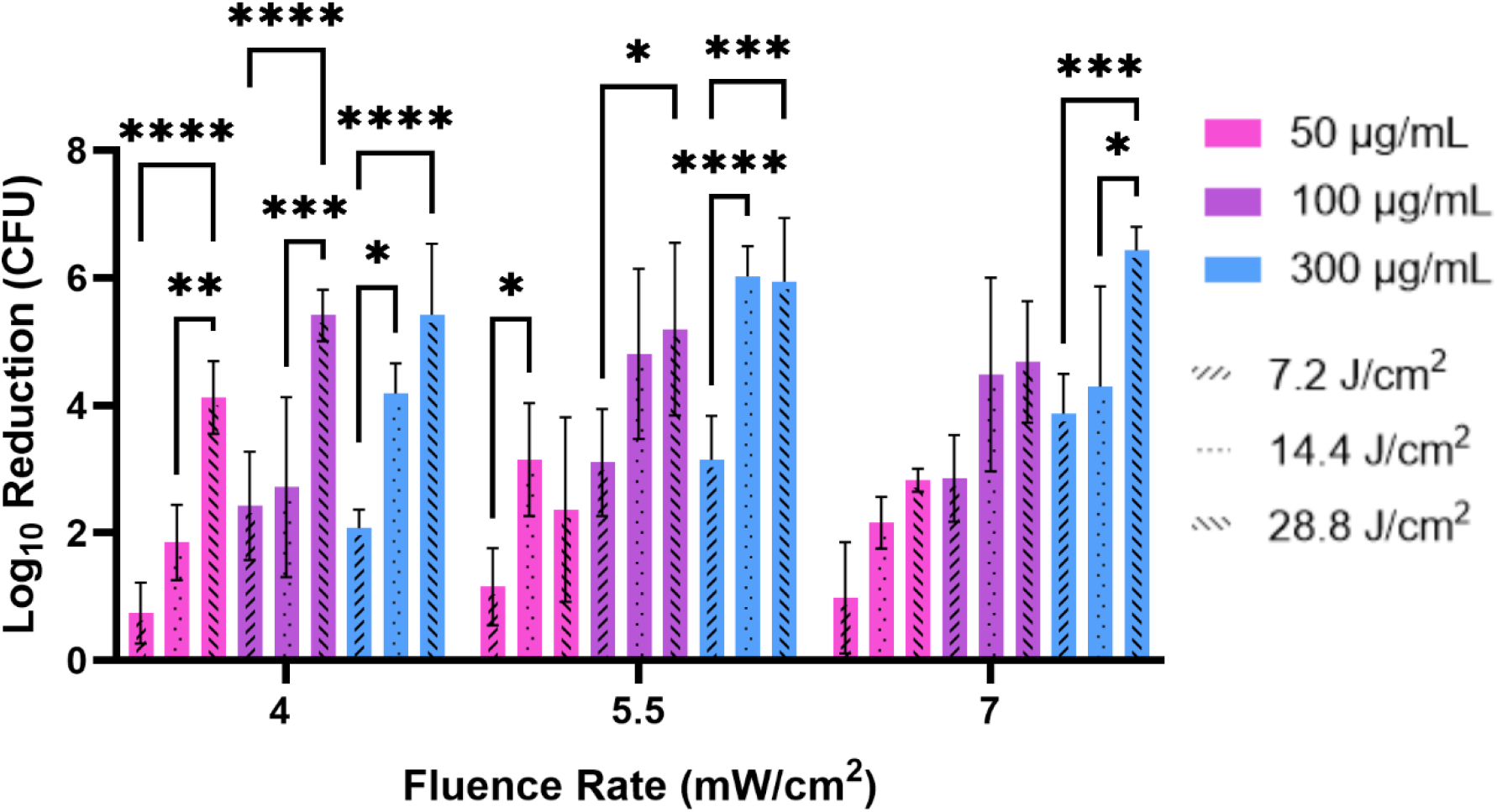
Effects of MB concentration, fluence (J/cm^2^), and fluence rate (mW/cm^2^) on PDT efficacy in *P. aeruginosa*. * p<0.05, ** p<0.01, *** p<0.001, and **** p<0.0001.

## Discussion

We found that MB-PDT resulted in a substantial antimicrobial effect against *E. coli, S. aureus, E. faecalis*, and *P. aeruginosa*. While Gram-positive bacteria were eradicated for all combinations of fluence rate, fluence, and MB concentration tested, efficacy in Gram-negatives increased at higher MB concentrations and fluences. MB uptake was higher for Gram-positives relative to Gram-negatives, and was substantially lower in *P. aeruginosa* compared to *E. coli*. This reduced MB uptake could be compensated for by increasing the delivered light fluence in *P. aeruginosa*, although increasing the light fluence rate did not appear to have an effect.

While application of antimicrobial PDT to planktonic bacteria has been widely explored^1^, many of these studies focused on higher light doses, both in terms of fluence rate and fluence. For example, Schastak *et al* utilized 1 W/cm^2^ illumination up to a maximum fluence of 100 J/cm^2^ in order to achieve ≥6 log reduction in Gram-positive and Gram-negative bacteria^26^.. Further, Piksa *et al* reported modes of 100 mW/cm^2^ and 30 J/cm^2^ across 163 papers describing antimicrobial MB-PDT using a laser source of illumination^27^. On the other hand, we focused on PDT with lower fluence rates and fluence, based upon success with these parameters for similar bacteria demonstrated by Hadaris *et al*^15^. This focus on the efficacy of lower light doses is crucial for application to complicated clinical cases such as deep abdominal abscesses. In simulation, we have found that treatment planning with a lower fluence rate target leads to improved patient eligibility for PDT^13,21^. It is therefore important to determine efficacy *in vitro* at these reduced light doses, as this would enable PDT application in a wider patient population.

Prior literature has also largely focused on single combinations of photosensitizer concentration and light fluence/fluence rate, whereas we explored the effects of fluence rate and fluence independently. In oncologic PDT, fluence rate has been shown to be a key factor in determination of PDT efficacy^18^, particularly for the photosensitizers HPPH^19^ and PpIX^28^. This topic has been less explored in the antimicrobial space, with limited reports in the literature for specific illumination devices^29,30^. While we did not find an overall fluence rate effect in the current study, we did find that increasing the light fluence increased efficacy for *P. aeruginosa*. The importance of fluence has also been demonstrated for other Gram-negative bacteria, such as *Klebsiella pneumoniae*^24^, as well as biofilms^31,32^.

Compared to eukaryotic cells, bacteria tend to have negatively-charged surfaces thus necessitating the use of cationic, water-soluble photosensitizers such as methylene blue to ensure bacterial uptake^33^. We demonstrated slightly reduced MB uptake by *S. aureus* and *E. faecalis* compared to *E. coli*. Despite this small reduction in MB uptake, efficacy was increased for the two Gram-positive species relative to *E. coli*. This difference in efficacy could potentially be explained by the presence of the lipopolysaccharide outer membrane in Gram-negatives, which can also contain multiple reactive oxygen species (ROS) scavenging and reducing defense mechamisms^34^. Further, Gram-positive and Gram-negative bacteria show differential sensitivity to various species of ROS^35^. As MB can produce multiple ROS species upon photoactivation^36^, the relative rates of production could contribute to these observed differences. For *P. aeruginosa*, we observed reduced efficacy relative to Gram-positive and *E. coli*. As it appears that this reduction in efficacy can be overcome by increasing the light fluence, while light fluence rate did not have an effect, it seems that the absolute concentration of ROS produced is more important than the rate of ROS production.

We acknowledge some limitations in the current study. While we sought to utilize the most common Gram-positive and Gram-negative organisms found in abscesses, these represent a sub-sample of the total microbial population found clinically^5,15^. Experimental replicates were also drawn from the same initial cultures. Conclusions may therefore be specific to the organisms studied and exact culture conditions. We also examined bacteria grown in planktonic culture, rather than as biofilms. Although it is unclear whether biofilms form within abscesses^37^, this represents an important target for future study due to reductions in PDT susceptibility associated with biofilms^1^. Due to experimental constraints, MB uptake data were extracted from different culture dishes than those used for PDT experiments. It is therefore possible that the data presented in Figure 4 may be culture specific.

## Conclusion

We have demonstrated PDT efficacy against four representative bacteria commonly found in human abscesses at low fluence rate and fluence. By increasing the fluence to a modest value of 28.8 J/cm^2^, high efficacy was also achieved in challenging Gram-negative organisms such as *P. aeruginosa*. As the microbial population present in a patient abscess is not known at the time of treatment, this implies that minimum fluence rate and fluence targets that are efficacious against the range of potential bacteria present should be delivered. Achieving these targets over the entire site of infection appears readily achievable using our previously developed treatment planning software^13,20,21^, making PDT an exciting clinical option for this patient population.

## Acknowledgements

The authors would like to thank Dr. Laurel Baglia for assistance in preparing photodynamic therapy protocols. This work was funded by grants EB029921 and AI178152 from the National Institutes of Health.

## Notes

### Competing Interest Statement

The authors have declared no competing interest.

